# Targeting fatty acid beta-oxidation impairs monocyte differentiation and prolongs heart allograft survival

**DOI:** 10.1101/2022.02.09.479789

**Authors:** Yuehui Zhu, Hao Dun, Li Ye, Yuriko Terada, Leah P. Shriver, Gary J. Patti, Daniel Kreisel, Andrew E. Gelman, Brian W. Wong

## Abstract

Monocytes play an important role in the regulation of alloimmune responses after heart transplantation (HTx). Recent studies have highlighted the importance of immunometabolism in the differentiation and function of myeloid cells. While the importance of glucose metabolism in monocyte differentiation and function has been reported, a role for fatty acid β-oxidation (FAO) has not been explored. Heterotopic HTx was performed using hearts from Balb/c donor mice implanted into C57Bl/6 recipient mice and treated with etomoxir (eto), an irreversible inhibitor of carnitine palmitoyltransferase 1 (Cpt1), a rate-limiting step of FAO, or vehicle control. FAO inhibition prolonged HTx survival, reduced early T cell infiltration/ activation and reduced dendritic cell (DC) and macrophage infiltration to heart allografts of eto-treated HTx recipients. ELISPOT demonstrated eto-treated HTx were less reactive to activated donor antigen presenting cells. FAO inhibition reduced monocyte-to-DC and monocyte-to-macrophage differentiation in vitro and in vivo. Further, FAO inhibition did not alter the survival of heart allografts when transplanted into *Ccr2*-deficient recipients, suggesting the effects of FAO inhibition on reduced immune cell infiltration and increased heart allograft survival were dependent on monocyte mobilization. Finally, we confirmed the importance of FAO on monocyte differentiation in vivo using conditional deletion of *Cpt1a*. Our findings demonstrate that targeting FAO attenuates alloimmunity after HTx, in part through impairing monocyte differentiation.

## Introduction

Heart transplantation (HTx) is the preferred treatment for numerous patients with end-stage heart failure, providing a median survival of approximately 12 years and a significant improvement in quality of life (1, 2). Calcineurin inhibitors (FK506; cyclosporine) remain the primary immunosuppressive agent, despite the advent of newer immunosuppressants such as mTOR inhibitors (rapamycin (sirolimus); everolimus) or co-stimulation blockers (anti-CTLA4-immunoglobulin; Belatacept). All these immunosuppressive agents (including the adjuvant therapies of corticosteroids and mycophenolic acid) mechanistically target T cell proliferation, and despite robust suppression of acute T cell responses that have led to dramatic reductions in acute rejection, chronic allograft rejection remains.

During acute inflammation, classical monocytes are mobilized from the bone marrow and migrate to sites of injury (3). Monocytes can respond to factors such as granulocyte-macrophage colony-stimulating factor (GM-CSF) or macrophage colony-stimulating factor (M-CSF), among other factors, to differentiate into monocyte-derived cells that have phenotypic similarities to dendritic cells (DCs) and macrophages (4-7). In acute kidney allograft rejection, monocyte infiltration has been quantitatively associated with renal dysfunction (8), and monocyte-mediated acute renal rejection has been reported after inhibition of T cell infiltration by Alemtuzumab (9). A higher number of CD16^+^ monocytes pre-transplant was associated with higher risk of acute rejection after kidney transplantation (10). The monocyte-macrophage lineage is central to allograft rejection, contributing to alloimmunity via multiple mechanisms including antigen processing/presentation, co-stimulation and proinflammatory cytokine production (11, 12). CD16^+^ monocyte infiltration and polarized M2 macrophages in endomyocardial biopsies have been associated with acute cellular rejection in human heart transplants (13). Monocyte-derived DCs have also been reported to impair early graft function in islet transplantation (14), and can promote cardiac allograft rejection (15).

The role of cellular metabolic pathways in immune cells, termed immunometabolism, has recently emerged as a central regulator of immune cell differentiation and function (16-22). To date, a role for fatty acid β-oxidation (FAO) has not been explored in monocyte differentiation. Much of our insight on the role of oxidative metabolism has been garnered through in vitro analysis of cells using extracellular flux analysis, which only provides an assessment of oxygen consumption and extracellular (media) acidification as a surrogate for lactate production based on substrate utilization in the presence and absence of metabolic inhibitors (16, 20, 23). In this study, we aimed to test the effect of pharmacological inhibition of FAO using etomoxir (eto), an irreversible inhibitor of carnitine palmitoyltransferase 1 (Cpt1), a rate-limiting step of FAO, in a murine model of acute heart allograft rejection. Further, we verified the importance of FAO in monocyte differentiation in vivo using conditional deletion of *Cpt1a*.

## Results

### Pharmacological inhibition of FAO extends HTx survival

To investigate the role of FAO in HTx rejection, we performed syngeneic (B6 → B6) and allogeneic (Balb → B6) murine heterotopic cardiac transplants. Pharmacological inhibition of FAO with eto starting on the day of transplantation significantly prolonged allograft survival compared with vehicle control (ctrl) (median graft survival: 13 vs 8 days in eto-vs ctrl-treated HTx (Figure 1A). Histological analysis at 4 days post-transplant revealed reduced immune cell infiltration in eto-treated HTx (Figure 1B). To further define the cellular infiltrate preceding acute heart allograft rejection, flow cytometric analysis was performed on day 4 post-transplant. This revealed a significant reduction in the abundance of T cells, activated T cells, CD4^+^ T cells, CD8^+^ T cells and Foxp3^+^CD4^+^ T cells consistent with the phenotype of regulatory CD4^+^ T cells (T_reg_), in transplanted hearts from eto-treated HTx compared with ctrl-treated HTx (Figure 1, C-G). We did not observe a difference in the ratio of CD4^+^ to CD8^+^ T cells in transplanted hearts between the two groups (Supplemental Figure 1A). We also observed a significant reduction in CD11b^+^CD64^+^CD24^−^ macrophages and CD11c^+^MHCII^+^CD64^−^ DCs in transplanted hearts from eto-treated HTx compared to ctrl-treated HTx (Figure 2, A and B). In line with this observation, allografts from eto-treated HTx had significant reductions in both recipient-derived macrophages and DCs (Figure 2, C and D). Analysis of FAO inhibition in syngrafts did not reveal any significant differences in infiltrating leukocytes relative to the syngrafts receiving vehicle control (Supplemental Figure 1, B-H).

**Figure 1:**
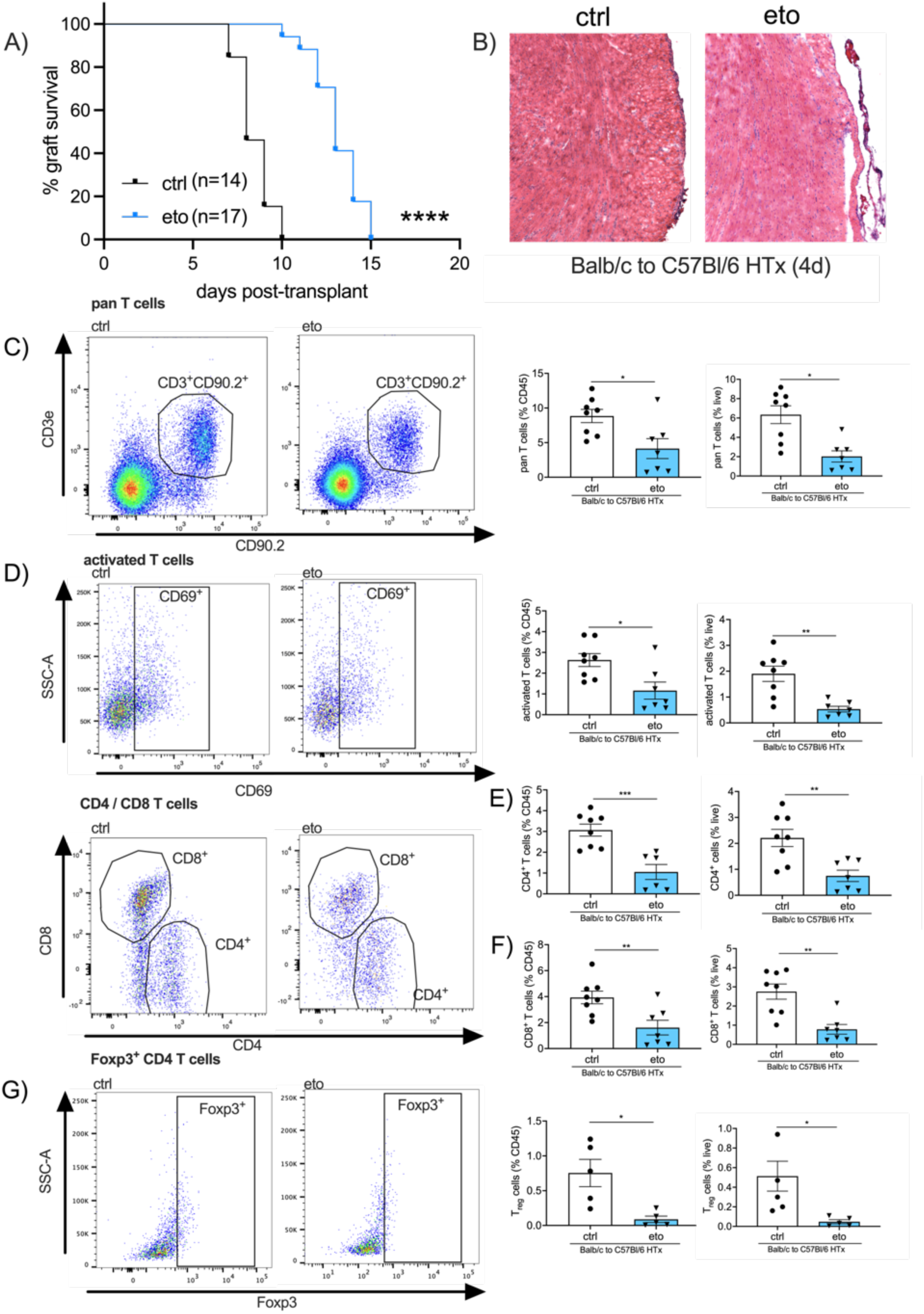
FAO inhibition improves HTx survival and decreases T cells in the transplanted heart. **A)** Comparison of graft survival in vehicle (ctrl)- or etomoxir (eto)-treated Balb → B6 heart allografts. The survival benefit was significant (\*\*\*\**p*<0.0001 by Mantel-Cox test). **B)** Representative micrographs at 4 days post-transplant showing reduced immune cell infiltration in eto-compared to ctrl-treated HTx captured at 20x magnification. **C-G)** Flow cytometric assessment in transplanted hearts from Balb → B6 heart allografts at 4 days post-transplant shown as a percentage of CD45^+^ cells (left graph) or live cells (right graph). **C)** Pan T cells assessed by CD3^+^CD90.2^+^ cells (ctrl, n=8; eto, n=7). **D)** Activated T cells assessed by CD69^+^CD3^+^CD90.2^+^ cells (ctrl, n=8; eto, n=7). **E)** CD4 T cells assessed by CD4^+^CD3^+^CD90^+^ cells (ctrl, n=8; eto, n=7). **F)** CD8 T cells assessed by CD8^+^CD3^+^CD90.2^+^ cells (ctrl, n=8; eto, n=7). **G)** T regulatory cells assessed by Foxp3^+^CD4^+^CD3^+^CD90.2^+^ cells (ctrl, n=5; eto, n=5). Data are represented as mean ± SEM. *, *p*<0.05; **, *p*<0.01; ***, p<0.001 by *t*-test.

**Figure 2:**
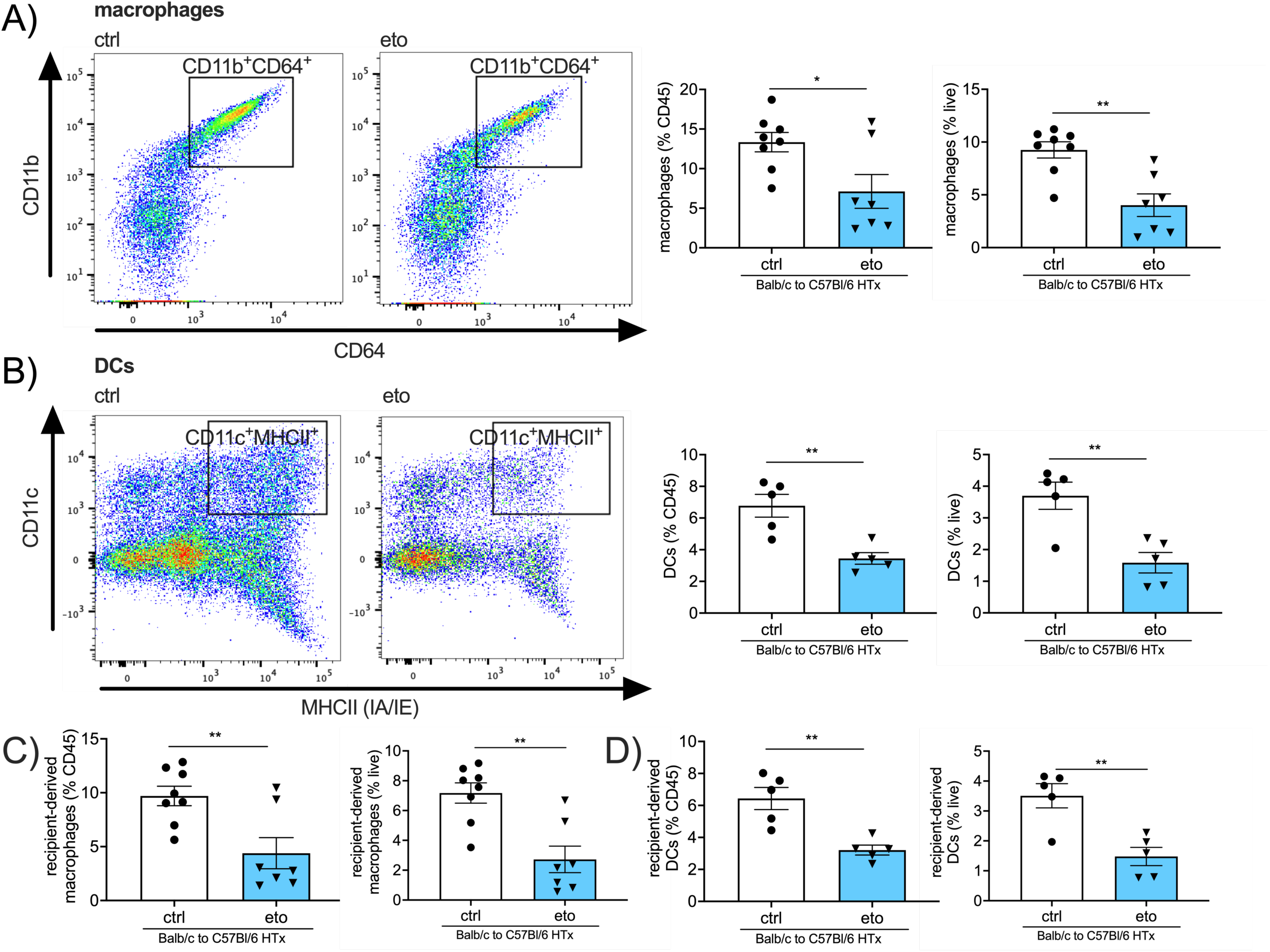
FAO inhibition reduces macrophages and DCs in the transplanted heart. **A-F)** Flow cytometric assessment in vehicle (ctrl)- or etomoxir (eto)-treated Balb → B6 heart allografts at 4 days post-transplant shown as a percentage of CD45^+^ cells (left graph) or live cells (right graph). **A)** Macrophages assessed by CD11b^+^CD64^+^CD24^−^ cells (ctrl, n=8; eto, n=7). **B)** Dendritic cells (DCs) assessed by CD11c^+^MHCII^+^ cells (ctrl, n=5; eto, n=5). **C)** Recipient-derived macrophages assessed by IA-b^+^CD11b^+^CD64^+^CD24^−^ cells (ctrl, n=8; eto, n=7). **D)** Recipient-derived DCs assessed by IA-b^+^CD11c^+^MHCII^+^ cells (ctrl, n=5; eto, n=5). Data are represented as mean ± SEM. *, *p*<0.05; **, *p*<0.01, by *t*-test.

### Inhibition of FAO reduces alloreactivity after HTx

We observed significant reduction in IFNγ, TNFα and IL2 production in splenocytes isolated from eto-treated HTx compared to ctrl-treated HTx recipients when stimulated with donor-, but not recipient-derived antigen presenting cells (APCs) (Figure 3, A-C; Supplemental Figure 2, A-C). In vitro inhibition of FAO in splenocytes from ctrl-treated HTx did not alter IFNγ production when stimulated with positive control (concanavalin A) or donor-derived APCs (Supplemental Figure 2, D and E), indicating that the effect of FAO in vivo is not likely due to direct T cell inhibition. Furthermore, FAO inhibition in splenocytes that were isolated from ctrl-treated HTx allograft recipients did not alter their proliferative responses after stimulation with donor-derived APCs (Supplemental Figure 2F).

**Figure 3:**
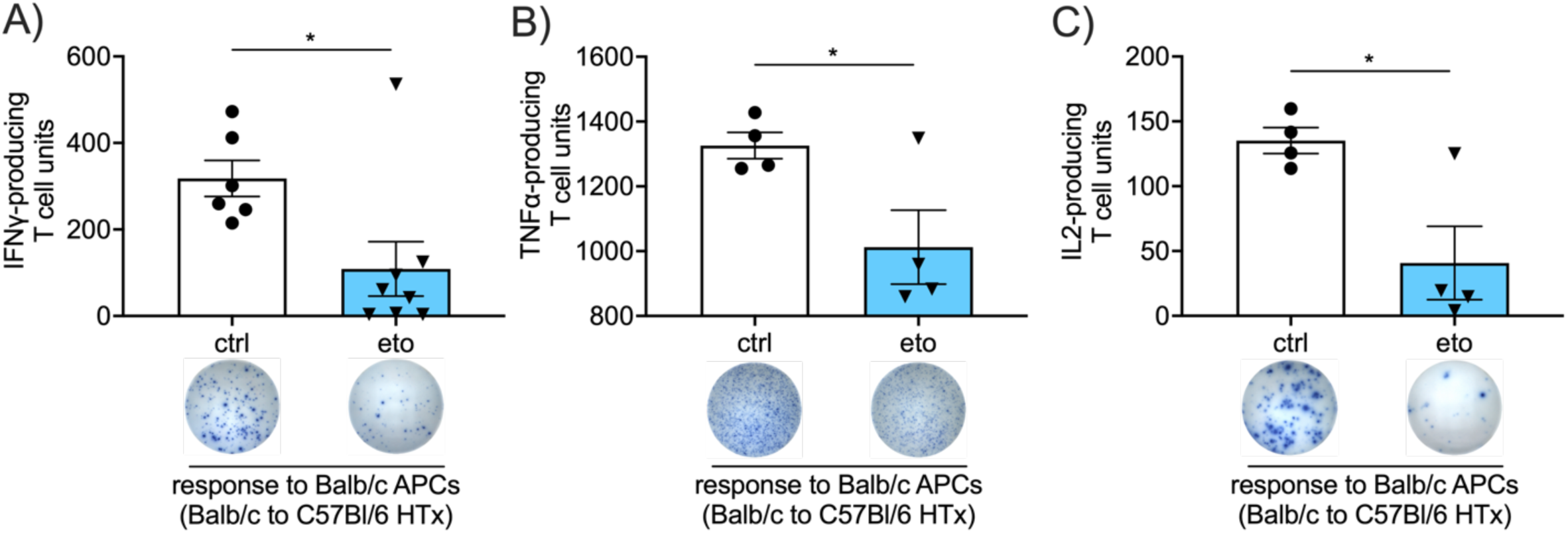
FAO inhibition reduces alloreactivity after HTx. **A-C)** ELISPOT assay performed using splenocytes procured at 4 days post-transplantation from vehicle (ctrl)- or etomoxir (eto)-treated Balb → B6 heart allografts. Samples were stimulated with donor-derived (Balb) APCs and assessed for the production of IFNγ (ctrl, n=6; eto, n=8) **(A)**, TNFα (ctrl, n=4; eto, n=4) **(B)** or IL2 (ctrl, n=4; eto, n=4) **(C)**. Data are represented as mean ± SEM. *, *p*<0.05 by *t*-test.

### FAO inhibition impairs monocyte differentiation after HTx

We next sought to investigate whether FAO plays a role in monocyte differentiation. To this end, we differentiated bone marrow-derived monocytes toward DC (GM-CSF + IL4) and macrophage (M-CSF) fates in the presence or absence of 100 µM eto for 7 days. Pharmacological inhibition of FAO significantly impaired the generation of cells with dendritic (Figure 4A) and macrophage (Figure 4B) phenotypes. Analysis of proliferation did not reveal any significant differences between eto- and ctrl-treated cells towards either fate (Supplemental Figure 3, A and B). Investigation of central carbon flux in monocytes showed a significant increase in FAO flux after 48 hours incubation following either GM-CSF + IL4 or M-CSF stimulation, which was sharply inhibited by pharmacological inhibition of FAO (Figure 4C). Inhibition of FAO flux in monocytes by eto was confirmed for a range of doses from 5-100 µM (Supplemental Figure 3C). Further, Cpt1 activity assay was used to verify the ability of eto to impair both palmitate and palmitoyl-CoA transport into isolated mitochondria from murine monocytes (Figure 4, D and E). Stimulation of monocytes with either GM-CSF + IL4 or M-CSF significantly increased both glycolytic flux (Supplemental Figure 3D) and cell proliferation (Supplemental Figure 3E). We did not observe any increased cell death due to inhibition of FAO, as assessed by LDH enzyme release, and FAO inhibition in GM-CSF + IL4-treated monocytes reduced LDH release, suggesting decreased cell death (Figure 4F).

**Figure 4:**
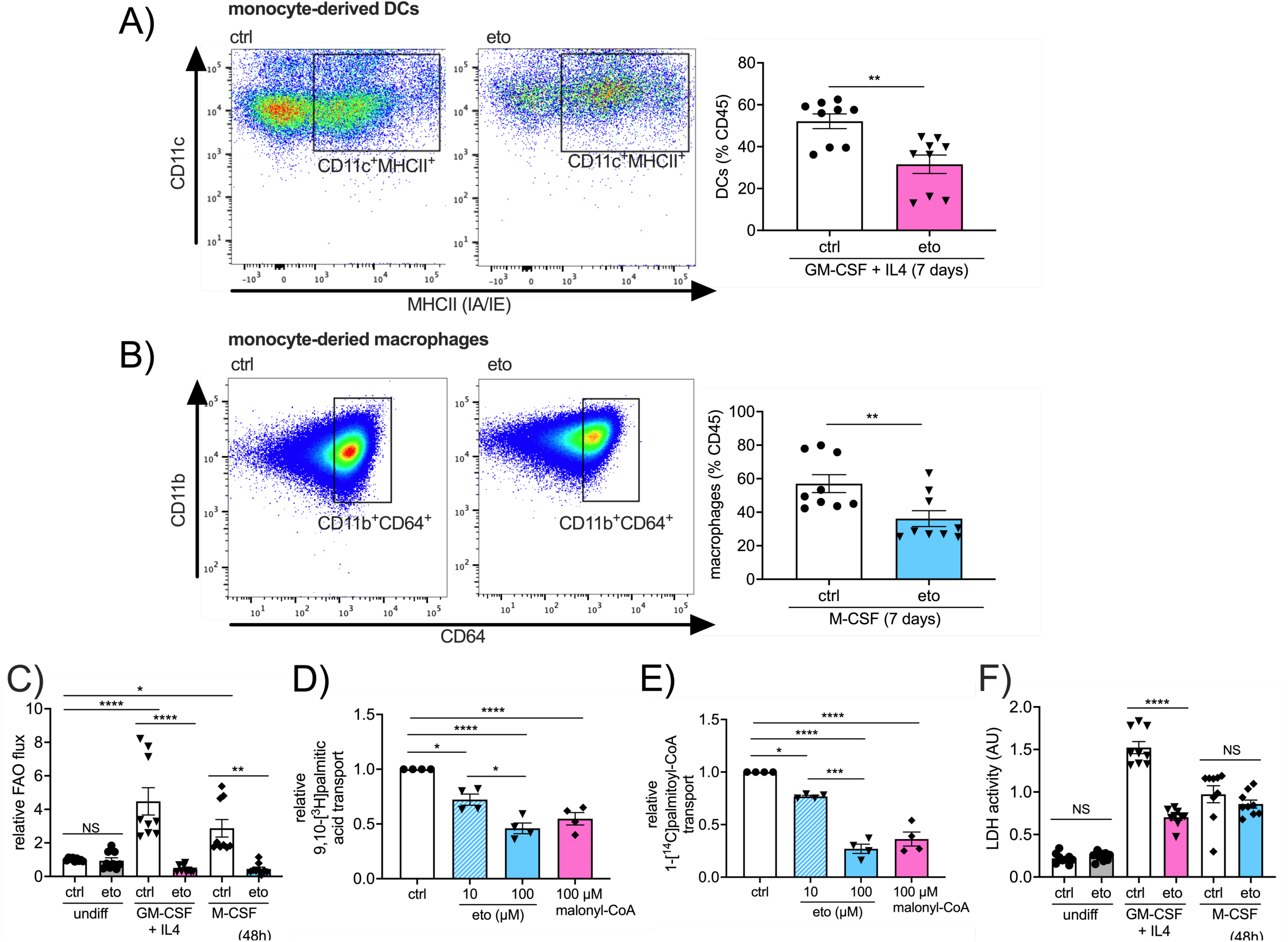
FAO inhibition impairs monocyte-to-DC and monocyte-to-macrophage differentiation in vitro. **A**,**B)** Flow cytometric assessment of monocytes after 7 days culture, shown as a percentage of CD45^+^ cells. **A)** Dendritic cells (DCs) assessed by CD11c^+^MHCII^+^CD64^−^ (ctrl, n=9; eto, n=9). **B)** Macrophages assessed by CD11b^+^CD64^+^CD24^−^ (ctrl, n=9; eto, n=9). **C)** FAO flux in monocytes in media only, or treated for 48 hours with GM-CSF + IL4 (DC differentiation condition) or M-CSF (macrophage differentiation condition), as assessed by [9,10-^3^H]palmitic acid radioisotopic incorporation (ctrl, n=9; eto, n=9). **D**,**E)** Cpt1 activity assay in intact mitochondria from monocytes treated with 10 or 100 µM etomoxir or 100 µM malonyl-CoA co-incubated with either 9,10-[^3^H]palmitic acid (**D**) (n=4) or 1-[^14^C]palmitoyl-€ (**E**) (n=4). **F)** Cell death in monocytes in media only, or treated for 48 hours with GM-CSF + IL4 or M-CSF, as assessed by lactate dehydrogenase (LDH) release (ctrl, n=9; eto, n=9). Data are represented as mean of individual data points from 3 independent experiments ± SEM. \**p*<0.05; \*\**p*<0.01; \*\*\**p*<0.001; \*\*\*\**p*<0.0001; NS, not statistically significant, by *t*-test **(A**,**B)** or ANOVA and Bonferroni post-hoc test **(C-F)**.

To investigate the role of FAO on monocyte differentiation in vivo, we adoptively transferred CD45.1^+^ monocytes at the time of heart transplantation into eto- or ctrl-treated recipients (Figure 5A). We observed significant reductions in CD45.1^+^ DCs and CD45.1^+^ macrophages in the allograft hearts of eto-treated recipients compared with ctrl-treated recipients (Figure 5, B and C). There were comparable abundances of Ki-67^+^ proliferating monocyte-derived cells in allografts between eto- and ctrl-treated recipients (Supplemental Figure 4, A and B), and no significant differences in the relative abundance of CD45.1^+^ cells in transplanted hearts, native heart, spleen or mediastinal lymph node (Supplemental Figure 4C). Moreover, the relative abundances of CD45.1^+^ DCs or CD45.1^+^ macrophages were comparable in the native hearts, spleen and mediastinal lymph nodes of eto- and ctrl-treated HTx recipients (Supplemental Figure 4, D and E). Analysis of CD45.2^+^ DCs and macrophages revealed a significant reduction in these populations in transplanted hearts from eto-treated HTx recipients compared with ctrl-treated HTx (Supplemental Figure 4, F and G).

**Figure 5:**
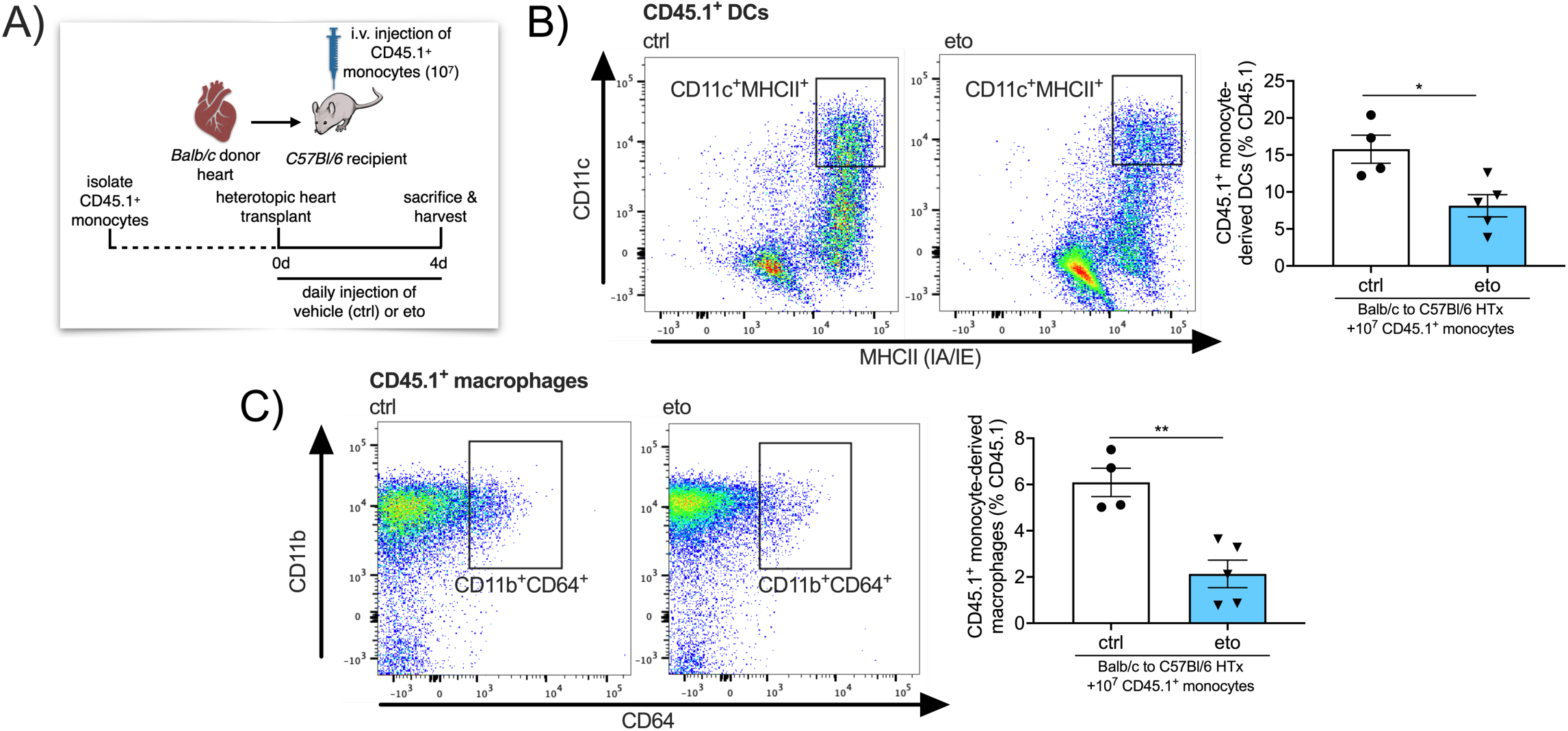
FAO inhibition impairs monocyte-derived DC and monocyte-derived macrophage differentiation in heart allografts in vivo. **A)** Schematic representation of adoptive transfer of 10^7^ CD45.1^+^ monocytes isolated from *Pepboy* mice intravenously injected into HTx recipients at the time of engraftment. **B**,**C)** Flow cytometric assessment of Balb → B6 heart allografts at 4 days post-HTx, adoptively transferred CD45.1^+^ monocytes. Cell subtypes are shown as a percentage of CD45.1^+^ cells. **B)** CD45.1^+^ DCs assessed by CD11c^+^MHCII^+^CD64^−^ cells (ctrl, n=4; eto, n=5). **C)** CD45.1^+^ macrophages assessed by CD11b^+^CD64^+^CD24^−^ cells (ctrl, n=4; eto, n=5). Data are represented as mean ± SEM. \**p*<0.05; \*\**p*<0.01; NS, not statistically significant, by *t*-test.

### HTx survival benefit from FAO inhibition is abrogated in *C**cr**2*^*-/-*^recipients

To further assess whether the HTx survival benefit observed from pharmacological inhibition of FAO was related to its effects on monocytes, we transplanted Balb hearts into B6 Ccr2-deficient (*Ccr2*^-/-^) recipients. We and others have shown that monocyte mobilization from the bone marrow is dependent on Ccr2 (24-26). Consistent with previous reports we observed a modest prolongation in allograft survival when recipients lack Ccr2 (27, 28). Importantly, heart allograft survival in *Ccr2*^*-/-*^recipients was independent of FAO inhibition (median HTx survival: 14 vs 14 days post-transplant in ctrl vs eto-treated *Ccr2*^*-/-*^HTx; Figure 6A). Of note, the median survival of hearts that were transplanted into either ctrl- and eto-treated *Ccr2*^*-/-*^HTx recipients was comparable to Balb hearts transplanted into wildtype B6 allogeneic hosts that were treated with eto. We observed similar relative abundances of total and activated T cells within transplanted hearts between ctrl- and eto-treated *Ccr2*^*-/-*^HTx recipients, which were also comparable to our observations in cardiac allografts after transplantation into eto-treated B6 recipients (Figure 6, B and C). Additionally, there were no significant differences in the relative abundances of DCs, macrophages, CD4^+^ or CD8^+^ T cells or T_reg_ cells in allografts in eto-versus ctrl-treated *Ccr2*^*-/-*^ HTx recipients, suggesting that the reduction in these immune cell populations induced by FAO inhibition in Balb to B6 HTx were secondary to the presence of Ccr2-positive cells in the transplant recipient (Figure 6, D and E; Supplemental Figure 5, A-C). As expected, heart allografts from *Ccr2*^*-/-*^HTx recipients had significantly reduced numbers of DCs compared with heart allografts from Balb to B6 HTx recipients, with a trend towards reduced numbers of macrophages (Figure 6, D and E). Further, investigation on recipient-derived populations of DCs and macrophages revealed a robust decrease in these populations in Balb to *Ccr2*^*-/-*^HTx recipients (Supplemental Figure 5, D and E). Finally, assessment of T cell alloreactivity showed no significant differences between splenocytes from ctrl- and eto-treated *Ccr2*^*-/-*^HTx when stimulated with donor- or recipient-derived APCs (Figure 6F; Supplemental Figure 5F).

**Figure 6:**
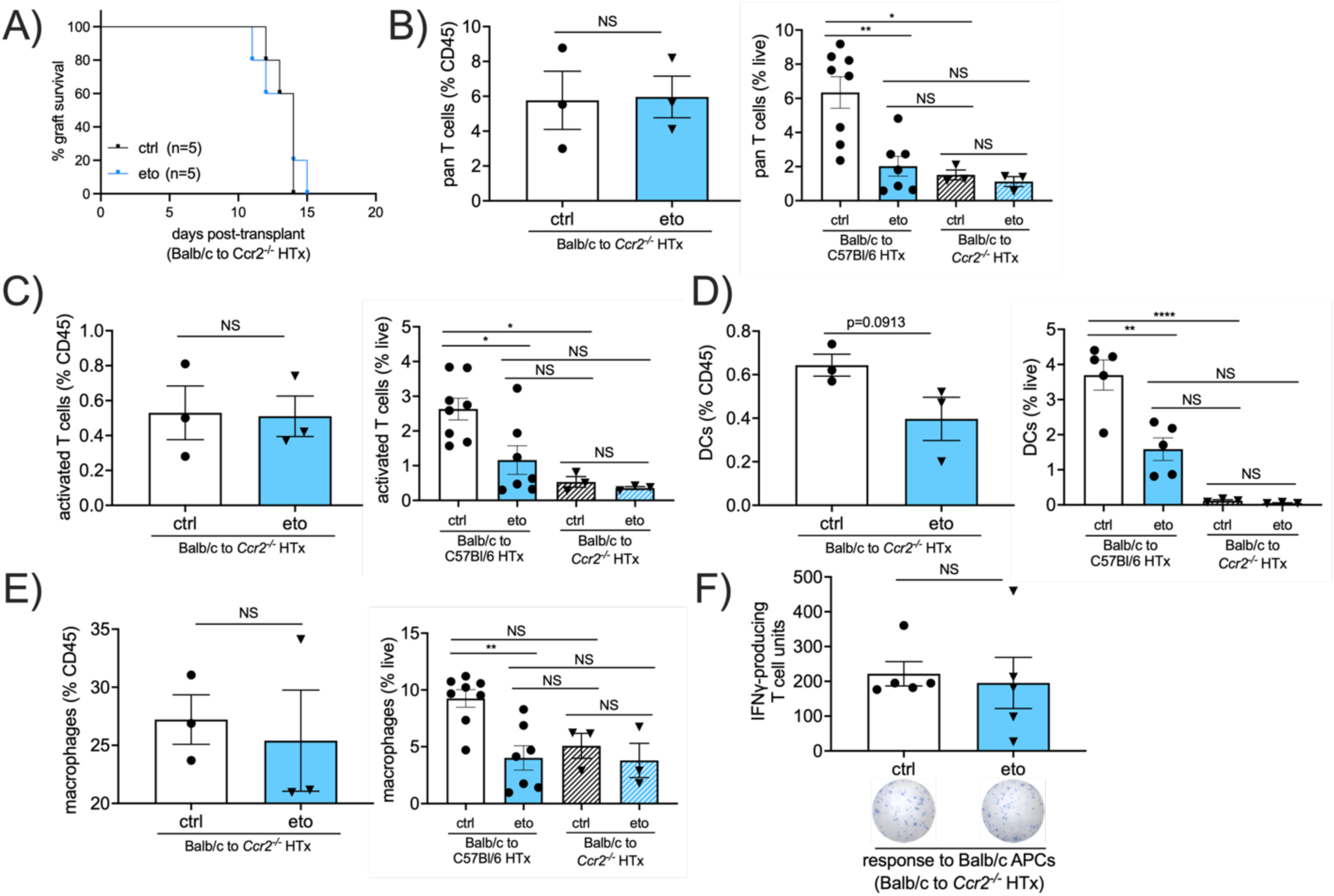
Loss of Ccr2 in HTx recipients abrogates the effects of FAO inhibition. **A)** Comparison of graft survival in vehicle (ctrl)- or etomoxir (eto)-treated Balb → B6 CCR2^-/-^heart allografts. There was no statistical difference in survival (p=0.7698 by Mantel-Cox test). **B-E)** Flow cytometric assessment in Balb → *Ccr2*^*-/-*^heart allografts at 4 days post-transplant shown as a percentage of CD45^+^ cells (left graph) or live cells (right graph; data for live cells from Balb → B6 HTx is re-plotted from Figs. 1 and 2). **B)** Pan T cells assessed by CD3^+^CD90.2^+^ (ctrl, n=8; eto, n=7 for Balb → B6 HTx; ctrl, n=3; eto, n=3 for Balb → B6 *Ccr2*^*-/-*^HTx). **C)** Activated T cells assessed by CD69^+^CD3^+^CD90.2^+^ cells (ctrl, n=8; eto, n=7 for Balb → B6 HTx; ctrl, n=3; eto, n=3 for Balb → B6 *Ccr2*^*-/-*^HTx). **D)** Dendritic cells (DCs) assessed by CD11c^+^MHCII^+^CD64^−^ cells (ctrl, n=5; eto, n=5 for Balb → B6 HTx; ctrl, n=3; eto, n=3 for Balb → B6 *Ccr2*^*-/-*^HTx). **E)** Macrophages assessed by CD11b^+^CD64^+^CD24^−^ cells (ctrl, n=8; eto, n=7 for Balb → B6 HTx; ctrl, n=3; eto, n=3 for Balb → B6 *Ccr2*^*-/-*^HTx). **F)** ELISPOT assay performed using splenocytes procured at 4 days post-transplantation from vehicle (ctrl) or etomoxir (eto)-treated Balb → B6 *Ccr2*^*-/-*^heart allografts. Samples were stimulated with donor-derived (Balb) APCs and assessed for the production of IFNγ (ctrl, n=5; eto, n=5). Data are represented as mean ± SEM. \**p*<0.05; \*\**p*<0.01; \*\*\*\**p*<0.0001; NS, not statistically significant by *t*-test (**B-E left graphs, F**) or ANOVA and Bonferroni post-hoc test (**B-E right graphs**).

### Genetic deletion of Cpt1a in monocytes impairs monocyte differentiation after HTx

To further confirm the importance of FAO in monocyte differentiation, we generated transgenic mice with conditional deletion of *Cpt1a* in monocytes (*Ccr2*.*Cre*^*ER*^*;Cpt1a*^*fl/fl*^, termed *Ccr2*^*ΔCpt1a*^). Littermate Cre-negative, *Cpt1a*^*fl/fl*^ mice were used as controls (termed wildtype). We adoptively transferred wildtype or *Ccr2*^*ΔCpt1a*^ monocytes at the time of transplantation into CD45.1^+^ Balb/c hearts to CD45.1^+^ B6 recipients (Figure 7A) and observed significant reductions in the relative abundance of CD45.2^+^ DCs and CD45.2^+^ macrophages in transplanted hearts from recipients adoptively transferred with *Ccr2*^*ΔCpt1a*^ compared to wildtype monocytes (Figure 7, B and C). In vivo deletion of Cpt1a in Ccr2-expressing cells did not alter monocyte numbers in the bone marrow (Supplemental Figure 6A). Assessing Ki-67^+^ proliferating cells from transplanted hearts in recipients adoptively transferred with wildtype or *Ccr2*^*ΔCpt1a*^ monocytes revealed no significant difference in *Ccr2*^*ΔCpt1a*^ monocyte-derived DCs or *Ccr2*^*ΔCpt1a*^ monocyte-derived macrophage Ki67 positivity compared with their wildtype counterparts (Supplemental Figure 6, B and C). Finally, we did not observe a difference in CD45.2^+^ cells in the transplanted hearts of HTx recipients adoptively transferred with either *Ccr2*^*ΔCpt1a*^ monocytes or wildtype monocytes (Supplemental Figure 6D).

**Figure 7:**
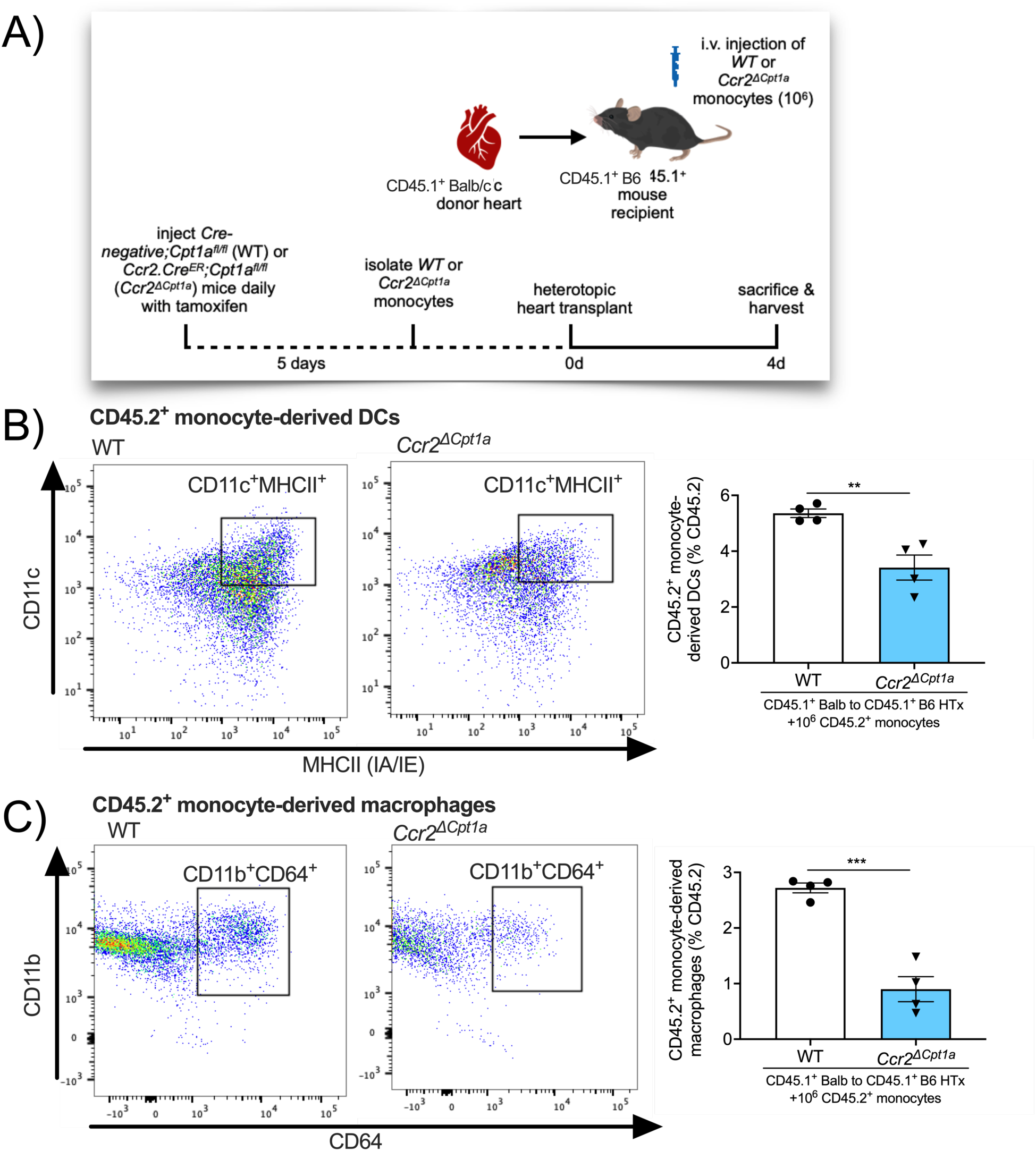
Genetic deletion of Cpt1a monocyte-derived DC and monocyte-derived macrophage differentiation in heart allografts in vivo. **A)** Schematic representation of adoptive transfer of 10^6^ monocytes isolated from *Cre-negative;Cpt1a*^*fl/fl*^ (WT) or *Ccr2*.*Cre*^*ER*^*;Cpt1a*^*fl/fl*^ (*Ccr2*^*ΔCpt1a*^) mice intravenously injected into CD45.1^+^ Balb/c to CD45.1^+^ B6 HTx recipients at the time of engraftment. **B-D)** Flow cytometric assessment of transplanted hearts from CD45.1^+^ Balb → CD45.1^+^ B6 allografts at 4 days post-HTx, adoptively transferred with WT or *Ccr2*^*ΔCpt1a*^ monocytes. All cell subtypes are shown as a percentage of CD45.2^+^ cells. CD45.2^+^ DCs assessed by CD11c^+^MHCII^+^CD64^−^ cells (WT, n=4; *Ccr2*^*ΔCpt1a*^, n=4). **C)** CD45.2^+^ macrophages assessed by CD11b^+^CD64^+^CD24^−^ cells (WT, n=4; *Ccr2*^*ΔCpt1a*^, n=4). Data are represented as mean ± SEM. \*\**p*<0.01; \*\*\**p*<0.001; NS, not statistically significant, by *t*-test.

## Discussion

In our current study, we demonstrate that pharmacological inhibition of FAO significantly prolongs heart allograft survival. Adoptive transfer experiments revealed that FAO inhibition results in a significant impairment in monocyte differentiation which results in a reduction of monocyte-derived DCs and macrophages in the transplanted heart, with no impairment on the homing of monocytes to the heart allograft. In vitro experiments confirmed that FAO inhibits monocyte differentiation, while not directly impacting T cell reactivity. Eto did not extend allograft survival in *Ccr2*^*-/-*^recipients, further supporting FAO-mediated dampening of alloimmunity through its effects on monocytes. Finally, genetic deletion of *Cpt1a* in monocytes mirrored effects on monocyte differentiation in heart allografts observed with pharmacological inhibition of FAO. Collectively, these findings present a previously unreported role for FAO in monocyte differentiation in the context of the regulation of alloimmunity.

Due to a reported role of glycolysis in macrophages and DCs, with sparse reports on specific oxidative metabolic pathways, we sought to investigate the role of FAO on heart allograft rejection. FAO has been reported to support an LPS-induced phenotype resembling the Warburg effect in glucose-deprived monocytes (23). However, this conclusion was primarily supported by results obtained by extracellular flux analysis. Raulien et al. reported that glutamine and fatty acids drive oxidative metabolism in the absence of glucose, and for FAO, this occurs at the expense of lipid droplets (23). While this altered metabolic state in glucose deprivation is reported to provide sufficient energy to sustain functional properties like cytokine secretion, migration and phagocytosis, it does not prevent a rise in AMP/ATP ratio or decreased respiratory burst (23). In the context of aging, monocytes are key producers of inflammatory cytokines, and monocytes from aged individuals were found to contain high levels of lipid droplets and impaired FAO, in part, through the downregulation of peroxisome proliferator-activated receptor-alpha (29), supporting the importance of FAO in monocyte function.

It is important to note that glycolysis is important in monocyte functions, including their activation and adhesion (30), and also plays a role in modulating CD8+ T cell function in human Chagas disease (31) and neutrophils in human hepatocellular carcinoma (32). Lee et al. demonstrated that LPS stimulation increases the rate of glycolysis in human classical monocytes, and that 2-deoxyglucose-mediated inhibition of glycolysis impaired LPS-induced activation and adhesion of monocytes (30). In this setting, increased glycolysis was regulated by mTOR-induced glucose transporter 1 expression, which subsequently led to increased accumulation of reactive oxygen species and activation of p38 MAPK, resulting in subsequent activation and adhesion of monocytes. However, these studies did not explore a direct role for FAO in monocyte function, in particular, in the context of an in vivo allogeneic environment.

As with all pharmacological approaches, FAO inhibition by eto would be expected to also affect other cell types that may be reliant on FAO, including cardiomyocytes (33), skeletal muscle cells (34), endothelial cells (35-37), and renal tubular epithelial cells (38). Within immune cell subsets, FAO has also been reported to be important in macrophage polarization (39) and CD8^+^ T memory cells (T_mem_) (40), however, recent evidence using genetic models has brought into question the specific requirement for Cpt1a or long chain FAO in M2 macrophage polarization (41) and T_mem_ cells (42, 43). Consistent with reports that Cpt1a/FAO is dispensable in T cells (42, 43), we did not observe direct effects of pharmacological inhibition of FAO on T cell activation. Eto has been shown to affect antigen-stimulated IL15 T_mem_-like cells in vitro (44), however, genetic studies targeting FAO enzymes revealed no impairment in T_mem_ cell formation in vivo (42). In our work, we did not profile for T_mem_ cells, however, the abrogation of survival benefit and T cell changes in *Ccr2*^*-/-*^recipient mice support the hypothesis of a FAO-dependent role in monocytes or monocyte-derived cells in our experiments. Administration of eto has been reported to improve cardiac function in fatty acid-induced ischemic injury (45), left ventricular performance in pressure-overloaded hearts (46-48), and also been tested in limited clinical trials in patients with congestive heart failure, demonstrating improved ejection fraction and cardiac output (49, 50). In our studies, we did not observe any further heart allograft survival benefit in *Ccr2*^*-/-*^HTx, suggesting that the efficacy of eto is dependent on the presence of Ccr2-expressing cells in the recipient. Of note, the mouse model of heterotopic heart transplantation results in a “non-functional” graft, where blood flows from the recipient aorta into the donor ascending aorta, supplying the coronary arteries. Blood then drains into the right atrium via the coronary sinus, which is then pumped into the right ventricle and subsequently enters the recipient inferior vena cava through the pulmonary artery. Coronary supply of arterial blood re-establishes sinus rhythm in the transplanted heart after reperfusion, however, the left chambers of the heart remain pressure under-loaded (51).

One limitation of the current study is our use of eto as a pharmacological inhibitor of FAO, which has been reported to have off-target effects affecting oxidative metabolism depending on dose and cell type (52, 53). We used this drug due to our extensive previous experience with its pharmacokinetics and pharmacodynamics in multiple cell types both in vitro and in vivo (35-37, 52, 53). Using a dose of 100 µM *in vitro*, we aimed to avoid some of the off-target effects affecting Complex I in the electron transport chain, or coenzyme A homeostasis described above 100 µM (specifically 200 µM and above) (41-43, 54). We further confirmed that eto was able to inhibit FAO flux and Cpt1 activity at multiple doses in murine monocytes. In our in vitro model of monocyte differentiation, we were able to demonstrate a significant reduction in FAO upon treatment with eto, however in vivo assessment of metabolic fluxes remains challenging. Off-target effects of eto, and the doses which elicit them, are harder to investigate. However, our experiments using *Ccr2*^*-/-*^recipient mice demonstrating the abrogation of the effect of pharmacological inhibition of FAO support our conclusion of the importance of FAO in monocyte differentiation. In addition, the role of FAO in monocyte differentiation in the context of heart transplantation in vivo was further verified using conditional deletion of *Cpt1a* in monocytes. The mechanism whereby FAO regulates monocyte differentiation remains to be determined. Our observation that both pharmacological inhibition of Cpt1, as well as genetic deletion of Cpt1a can both reduce monocyte-to-macrophage and monocyte-to-DC differentiation in heart allografts support a role for FAO in these functions. Further, investigation into the specific roles of FAO in DCs and macrophages in allograft rejection are also required. In addition to promoting alloimmunity and allograft rejection, monocytic populations may also play an important role in limiting alloimmune responses. It has previously been demonstrated that CD11b^+^CD115^+^Ly6C^+^ monocytes are required to maintain heart transplant tolerance induced by donor splenocyte transfusion in addition to anti-CD40L monoclonal antibody (55). Early after transplantation, monocytes migrate from the bone marrow to the transplanted organ, where they can prevent the initiation of adaptive immune responses that lead to allograft rejection and participate in the development of T_reg_ cells. It has been reported that reducing monocyte-derived macrophages impairs transplant tolerance, in part through increasing both CD8^+^ T cell numbers and reducing T_reg_ expansion (56, 57), suggesting that the role monocytes play in allogeneic immune responses may vary between acute and tolerogenic settings. As our work demonstrates that inhibition of FAO can impair the differentiation of monocyte-derived cells in acute HTx rejection, it would be of interest to further explore the role of FAO in monocyte dynamics in tolerogenic settings. While the potential clinical utilization of eto is unlikely, as clinical trials for eto were halted in phase II due to unacceptably high liver transaminase levels and associated hepatotoxicity (58, 59). Nevertheless, our study highlights the potential clinical utility of other pharmacological inhibitors of FAO, which warrants further investigation.

## Methods

### Animals

6-10 week old Balb/c (Balb) mice (#000651), C57Bl/6 (B6) mice (#000664), *B6*.*SJL-Ptprc*^*a*^*Pepc*^*b*^*/BoyJ* (known as *Pepboy*, CD45.1^+^ B6 background; #002014), *CByJ*.*SJL(B6)-Ptprc*^*a*^*/J* (CD45.1^+^ Balb background; #006584), *B6*.*129S4-Ccr2*^*tm1lfc*^*/J* (Ccr2-deficient mice; known as *Ccr2*^*-/-*^; #004999) and *B6(Cg)-Cpt1atm1c(EUCOMM)Hmgu/RjnsJ* (*Cpt1a*^*fl/fl*^ mice (36, 42); #032778) were purchased from Jackson Laboratory. *Ccr2*.*Cre*^*ER*^ transgenic mice were a generous gift from Dr. Andrew Gelman (WUSM). *Ccr2*.*Cre*^*ER*^ mice were intercrossed with *Cpt1a*^*fl/fl*^ mice to generate *Ccr2*.*Cre*^*ER*^*;Cpt1a*^*fl/fl*^ mice (termed *Ccr2*^*ΔCpt1a*^). Littermate Cre-negative *Cpt1a*^*fl/fl*^ mice were used as controls (termed wildtype (WT)). Animals were bred and housed under standard conditions.

### Reagents

Fluorochrome-conjugated antibodies used in this study are listed in Supplemental Table 1. LIVE/DEAD fixable violet dead cell stain kit was also from Thermo Scientific. Brilliant Stain Buffer Plus 1000t and BD Horizon Fixable Viability Stain 510 were from BD Biosciences. Concanavalin A, sodium hydroxide, trichloroacetic acid, (+)-etomoxir sodium salt hydrate, tamoxifen and other standard laboratory chemicals were from Sigma-Aldrich.

### Heterotopic heart transplant

Allogeneic (Balb to B6) and syngeneic (B6 to B6) heterotopic heart transplantation was performed as previously described (60, 61). Anesthesia was induced by a mixture of ketamine (80-100 mg/kg)/xylazine HCl (8-12 mg/kg), intraperitoneally (i.p.) and maintained with 1-2% isoflurane gas, as required. Briefly, donor hearts were implanted into the abdominal cavity of recipient mice, where the donor ascending aorta and pulmonary artery were anastomosed to the recipient infrarenal aorta and inferior vena cava, respectively. Donor hearts were maintained on ice between procurement and implantation, and cold ischemic times were less than 60 minutes. Heart transplant recipients were administered either vehicle (H_2_O) or etomoxir (20 mg/kg) daily by i.p. injection. Function of transplanted hearts was assessed by daily direct palpation along the abdomen adjacent to the implantation site until cessation of palpable heartbeat. Heart graft function was assessed using the following scale: +++, soft graft with strong contraction; ++, mild turgor and mild contraction; +, hard turgor and weak contraction; –, no palpable contraction. Cessation of palpable heartbeat was verified at time of sacrifice by laparotomy. For experiments involving *Ccr2*^*-/-*^mice, Balb donor hearts were transplanted into *Ccr2*^*-/-*^recipient mice. For experiments involving adoptive transfer of WT or *Ccr2*^*ΔCpt1a*^ monocytes, CD45.1^+^ Balb donor hearts were transplanted into CD45.1^+^ B6 recipient mice.

### Monocyte isolation, adoptive transfer of monocytes and in vitro differentiation

Bone marrow-derived monocytes were obtained from long bones using the MACS Monocyte Isolation Kit (Miltenyi Biotec) following manufacturer’s protocol. For adoptive transfer experiments, 10^7^ B6 CD45.1, monocytes were intravenously injected at time of HTx (62). For monocyte differentiation assays, 10^6^ monocytes were cultured in RPMI complete media (Invitrogen) supplemented with 10% FBS, penicillin/streptomycin (Invitrogen; 120 units/mL and 100 mcg/mL, respectively) and 0.05 mM 2-mercaptoethanol. Stimulation of monocytes to differentiate towards a DC or macrophage lineage was achieved by culturing with GM-CSF (25 ng/mL, Miltenyi) + IL-4 (25 ng/mL, Miltenyi) or M-CSF (100 ng/mL, Miltenyi), respectively. Monocytes were treated with 100 µM etomoxir for indicated experiments.

### Flow cytometry

Spleen and mediastinal lymph nodes were procured, rinsed with PBS, then the tissue was minced and digested with washing with 0.5 mg/mL collagenase I (Sigma) + 60 units/mL DNase I (Sigma) in DMEM, centrifuged and filtered through a 70 µm filter, then RBC lysis buffer (Sigma) was used, followed by a final centrifugation step and freezing using CTL-Cryo ABC Freezing Kit (CTL). Hearts were procured, perfused with cold PBS, then minced and digested with 450 units/mL collagenase I (Sigma) + 60 units hyaluronidase type 1-s (Sigma) + 60 units/mL Dnase I in DMEM, then centrifuged and filtered through a 40 µm filter, followed by RBC lysis and a final centrifugation step and freezing using CTL-Cryo ABC Freezing Kit (CTL). Flow cytometry was performed as previously described (63) using either a BD LSR Fortessa or a BD X-20 BD (BD Biosciences). Representative flow plots in the Figures illustrate the distribution of the target population, which is then presented in the associated graph(s) relative to CD45^+^ or live cells, unless otherwise indicated. Flow cytometry data was analyzed by FlowJo software (version 10.71).

For identification of total T cells, we used the following gating strategy: FSC/SSC, single live cells, CD45^+^, CD3^+^CD90.2^+^, CD69^+^, CD4^+^, CD8^+^ and Foxp3^+^CD4^+^ populations were identified from total T cell pools. For identification of DCs, we used the following gating strategy: FSC/SSC, live cells, CD45^+^, CD11c^+^MHCII^+^. For identification of macrophages, we used the following gating strategy: FSC/SSC, single live cells, CD45^+^, CD24^−^, CD11b^+^CD64^+^.

For adoptive transfer experiments, the following gating strategy was used for CD45.1^+^ DCs: FSC/SSC, single live cells, CD45^+^, CD45.1^+^, CD64^−^, CD11c^+^MHCII^+^ and the following gating strategy for macrophages: FSC/SSC, single live cells, CD45^+^, CD45.1^+^, CD24^−^, CD64^+^CD11b^+^. Similar gating strategies were used for the assessment of in vitro monocyte differentiation.

For experiments involving adoptive transfer of WT or *Ccr2*^*ΔCpt1a*^ monocytes, the following gating strategy was used for CD45.2^+^ DCs: FSC/SSC, single live cells, CD45^+^, CD45.2^+^, CD64^−^, CD11c^+^MHCII^+^ and the following gating strategy for CD45.2^+^ macrophages: FSC/SSC, single live cells, CD45^+^, CD45.2^+^, CD24^−^, CD11b^+^CD64^+^.

### ELISPOT

Pre-coated strips with antibodies directed against mouse interferon gamma (IFNγ), tumor necrosis factor (TNFα) or interleukin (IL)-2 (IL2) were purchased from Cellular Technology Limited (CTL). Splenocytes were obtained from HTx recipients, rinsed with PBS, then the tissue was minced and digested with 0.5 mg/mL collagenase I (Sigma) + 60 units/mL DNase I (Sigma) in DMEM, centrifuged and filtered through a 70 µm filter, then RBC lysis buffer (Sigma) was used, followed by a final centrifugation step and freezing using CTL-Cryo ABC Freezing Kit (CTL). Media only and 5 µg/mL of concanavalin A (conA; Sigma) were used as negative and positive controls, respectively. Isolated splenocytes from Balb or B6 mice were irradiated with 20 Gy using an X-RAD 320 Biological Irradiator (Precision X-Ray) and used as “activated” antigen-presenting cells (APCs). Splenocytes (100,000 cells) were co-incubated with positive control, negative control or activated APCs (100,000 cells) and cultured in CTL-test serum-free media (CTL) supplemented with penicillin/streptomycin (Invitrogen; 120 units/mL and 100 mcg/mL, respectively) and 2 mM L-glutamine (Invitrogen). 2-3 technical replicate wells were used for each condition, and n numbers represent samples from individual HTx. Single color ELISPOT assays were performed following manufacturer’s specifications, and plates were dried, scanned and analyzed using an Immunospot S6 Universal Analyzer (CTL).

### Histology

Tissues were procured and fixed in 4% paraformaldehyde before embedding in OCT for cryosections or processed for paraffin embedding. 5 µm sections were stained with hematoxylin and eosin to assess histopathological changes.

### Radioisotope tracer experiments

50,000 cells per well were seeded and allowed to attach before gentle washing and incubation for 48 hours in media containing 0.4 µCi/mL (24.24 nM final concentration) [5-^3^H]glucose (Perkin Elmer) or 2 µCi/mL (66.67 nM final concentration) [9,10-^3^H]palmitate (Perkin Elmer) for assessment of glycolytic or FAO flux, respectively. For FAO flux assays, 50 µM of L-carnitine solution (Sigma) was supplemented to the media. To assess the production of ^3^H_2_O after a cumulative 48 hours of drug treatment, media was transferred to glass vials and sealed with rubber stoppers. ^3^H_2_O was captured in hanging wells within the glass vials containing Whatman filter paper soaked with equivolume H_2_O over a period of 48 hours at 37 °C to reach saturation. Radioactivity for radioisotope tracer studies was determined by liquid scintillation counting using a Beckman Coulter LS6500 scintillation counter (Beckman Coulter, Inc.).

### Cpt1 activity assay

Cpt1 activity was performed on isolated mitochondria from murine monocytes Briefly, bone marrow-derived monocytes were obtained from long bones using the MACS Monocyte Isolation Kit (Miltenyi) following manufacturer’s protocol, then mitochondria were isolated using a mitochondria isolation kit (Thermo). Isolated mitochondria were co-incubated with radioisotope-labeled 9,10-[^3^H]palmitic and 1-[^14^C]palmitoyl-CoA in an assay solution containing 20 mM HEPES, 75 mM KCl, 2 mM KCN, 1% fatty acid-free BSA and 0.25 mM L-carnitine, adapted from previous publications (64-66). After incubating for 3 minutes at 37°C, the reaction was quenched with 3 M trichloroacetic acid (TCA), spun down at 12,000 x g for 5 minutes, then washed with 2 mM TCA and centrifuged again before removing the supernatant and extracting the mitochondrial pellet for liquid scintillation counting using a Beckman Coulter LS6500 scintillation counter (Beckman Coulter). The endogenous Cpt1 inhibitor malonyl-CoA (Sigma) was used as a positive control.

### Proliferation assay

Thymidine incorporation assay was used to assess cell proliferation. Briefly, 10,000 cells per well were seeded and incubated with 1 µCi/mL [^3^H]thymidine (Perkin Elmer) for specified time periods. Cells were fixed with 100% ethanol for 15 minutes at 4 °C, then precipitated with 10% trichloroacetic acid (Sigma) and lysed with 0.1N NaOH (Sigma). The amount of [^3^H]thymidine incorporated into DNA was measured by liquid scintillation counting. T cells were isolated from the spleens of HTx recipients using the pan T cell isolation kit (Miltenyi), then cultured in RPMI (Invitrogen) supplemented with 10% FBS, penicillin/streptomycin (Invitrogen; 120 units/mL and 100 mcg/mL, respectively), 0.05 mM 2-mercaptoethanol and 20 units/mL recombinant IL2 (Miltenyi). Monocytes were cultured as indicated above. T cells or monocytes were treated with 100 µM etomoxir for indicated experiments.

### Cytotoxicity assessment

Cytotoxicity Detection Kit (Roche CustomBiotech), which quantifies cytotoxicity/cytolysis based on the measurement of lactate dehydrogenase (LDH) release from damaged cells, was used to assess the effects of etomoxir on monocyte viability, as per manufacturer’s protocol.

### Statistics

All data are expressed as mean ± SEM, unless otherwise indicated. Student’s t test was used for two-group comparison. For multiple group analysis, ANOVA with Bonferroni post-hoc test was used to calculate statistical significance. Log-rank test (Mantel-Cox) test was used for survival curve comparison. P values less than 0.05 were considered statistically significant. Statistical analysis and graphics were performed using GraphPad Prism software and Microsoft Excel.

### Study approval

Animal experiments were conducted in accordance with an approved Washington University School of Medicine (WUSM) Institutional Animal Care and Use Committee (IACUC) protocol (#20190173; #20190174).

## Author contributions

AEG, DK and BWW designed the research studies. YZ, HD, LY, YT, BWW performed the experiments, including data acquisition. YZ and BWW analyzed the data. LS, GJP, DK and AEG assisted in interpretation of the data and critically reviewed the manuscript. BWW wrote the manuscript. The order of the co-first authors in the author list was decided as follows: YZ performed the flow cytometry, ELISPOT and in vitro experiments and HD established the heterotopic heart transplant protocol in mice in our laboratory and performed all the microsurgeries for this study, monitored transplanted heart function and health of animals throughout the course of experiments, administered drugs daily and harvested all tissues for downstream analysis. All authors read and discussed manuscript drafts.

## Acknowledgements

This work was supported by a grant from the American Society of Transplantation (AST) Research Network (BWW) and by the Joel D Cooper Career Development Award from the International Society for Heart and Lung Transplantation (ISHLT; BWW). This work was also supported by the Washington University Institute of Clinical and Translational Sciences which is, in part, supported by the NIH/National Center for Advancing Translational Sciences (NCATS), CTSA grant #UL1 TR002345. We acknowledge the assistance of the Digestive Disease Research Core Center (DDRCC) for histology, the assistance of the Bursky Center for Human Immunology & Immunotherapy Programs (CHiiPs) Core Laboratory for Immunospot assays, and the Flow Cytometry & Fluorescence Activated Cell Sorting Core in the Department of Pathology & Immunology and the Siteman Flow Cytometry Core at the Washington University School of Medicine for assistance with flow cytometry.

## Figures and Figure Legends

**Supplemental Figure 1:**
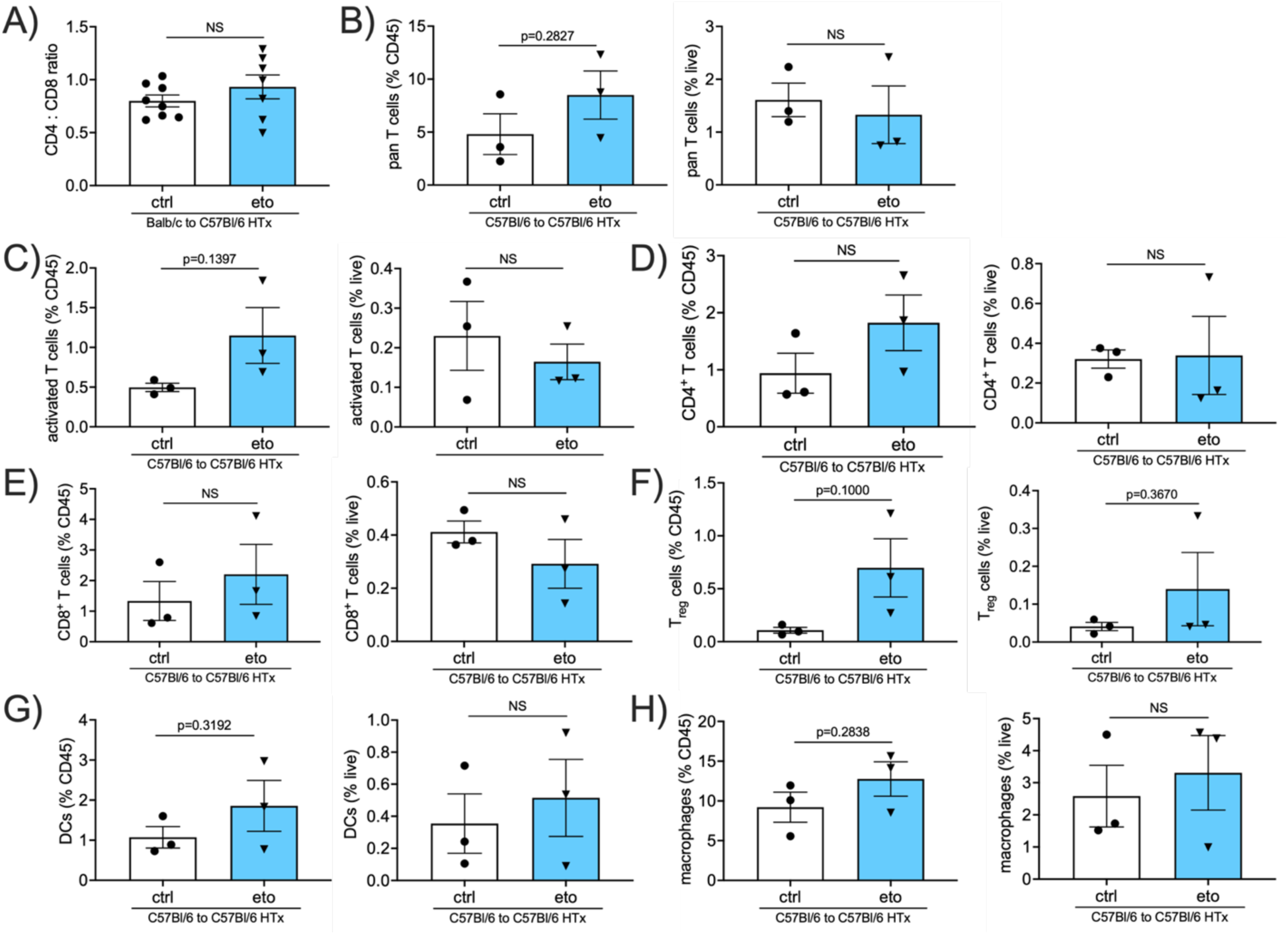
CD4:CD8 T cell ratio in HTx allografts and immune cell profile in HTx syngrafts. **A-H)** Flow cytometric assessment in transplanted hearts from Balb → B6 heart allografts at 4 days post-transplant shown as a percentage of CD45^+^ cells (left graph) or live cells (right graph). **A)** CD4^+^ to CD8^+^ T cell ratio from Balb → B6 heart allografts at 4 days post-transplant treated with vehicle (ctrl) or etomoxir (eto) (ctrl, n=8; eto, n=7). **B-H)** Flow cytometric assessment in transplanted hearts from B6 → B6 heart allografts at 4 days post-transplant shown as a percentage of CD45^+^ cells. **B)** Pan T cells assessed by CD3^+^CD90.2^+^ cells (ctrl, n=3; eto, n=3). **C)** Activated T cells assessed by CD69^+^CD3^+^CD90.2^+^ cells (ctrl, n=3; eto, n=3). **D)** CD4 T cells assessed by CD4^+^CD3^+^CD90.2^+^ cells (ctrl, n=3; eto, n=3). **E)** CD8 T cells assessed by CD8^+^CD3^+^CD90.2^+^ cells (ctrl, n=3; eto, n=3). **F)** T regulatory cells assessed by Foxp3^+^CD4^+^CD3^+^CD90.2^+^ cells (ctrl, n=3; eto, n=3). **G)** Dendritic cells (DCs) assessed by CD11c^+^MHCII^+^CD64^−^ cells (ctrl, n=3; eto, n=3). **H)** Macrophages assessed by CD11b^+^CD64^+^CD24^−^ cells (ctrl, n=3; eto, n=3). Data are represented as mean ± SEM. NS, not statistically significant by *t*-test.

**Supplemental Figure 2:**
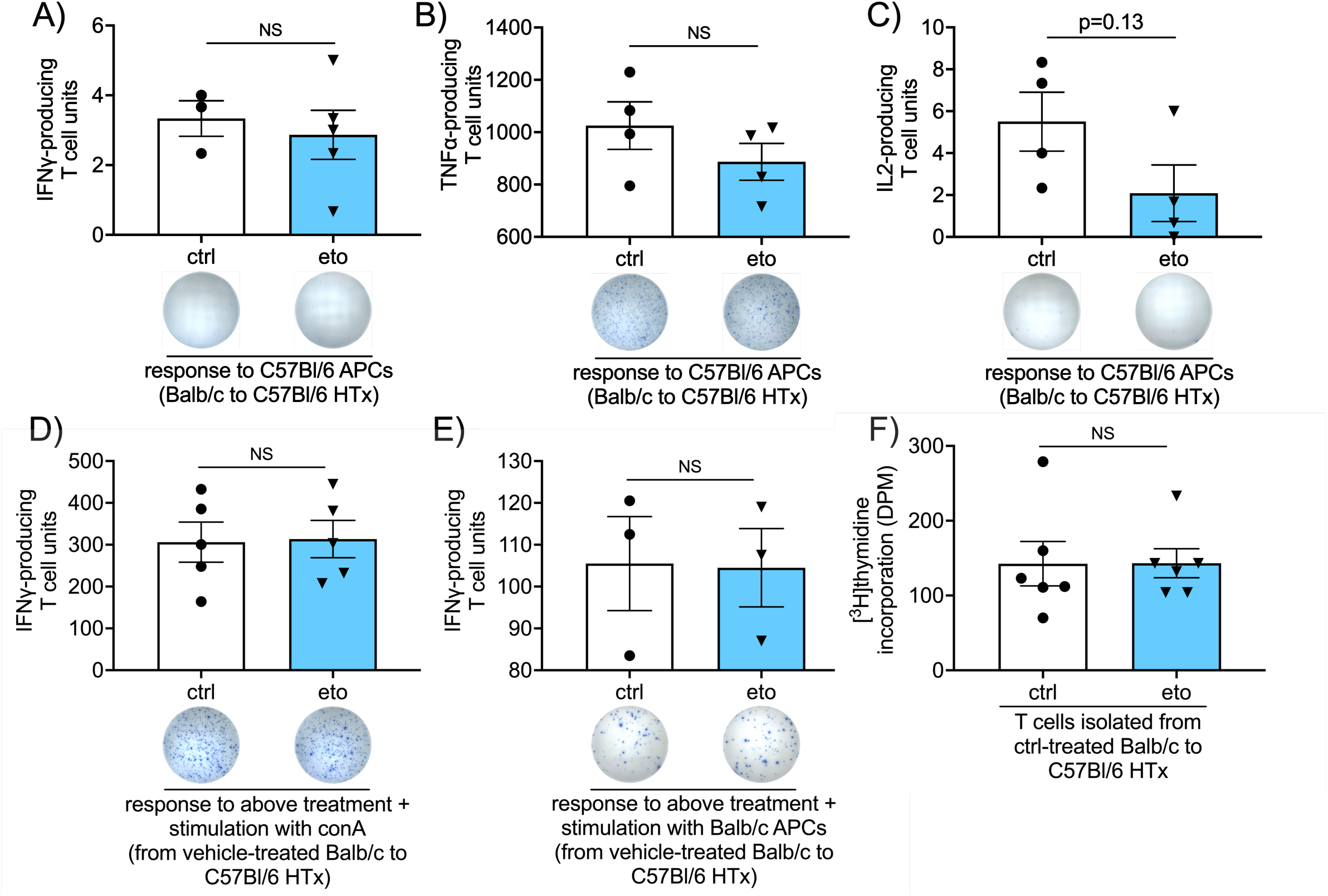
T cell activation in response to B6 APCs, direct effect of FAO inhibition on T cell activation and effect of FAO inhibition on T cell proliferation. **A-C)** ELISPOT assay performed using splenocytes procured at 4d post-transplantation from Balb → B6 heart allografts from either vehicle (ctrl) or etomoxir (eto)-treated animals. Samples were stimulated with activated B6 APCs and assessed for the production of IFNγ (ctrl, n=6; eto, n=8) (**A**), TNFα (ctrl, n=4; eto, n=4) (**B**) or IL2 (ctrl, n=4; eto, n=4) (**C**). **D**,**E)** ELISPOT assay performed using splenocytes procured at 4d post-transplantation from Balb → B6 heart allografts from ctrl-treated HTx and incubated in the presence or absence of eto. **D)** IFNγ production in response to positive control stimulus (concanavalin A (conA)) (ctrl, n=3; eto, n=3). **E)** IFNγ production in response to allostimulus (Balb APCs) (ctrl, n=3; eto, n=3). **F)** Proliferation of T cells isolated from spleens from ctrl-treated Balb → B6 HTx recipients in response to treatment with ctrl or eto for 24h, as assessed by [^3^H]thymidine incorporation (ctrl, n=6; eto, n=6). Data are represented as mean ± SEM. NS, not statistically significant by *t*-test.

**Supplemental Figure 3:**
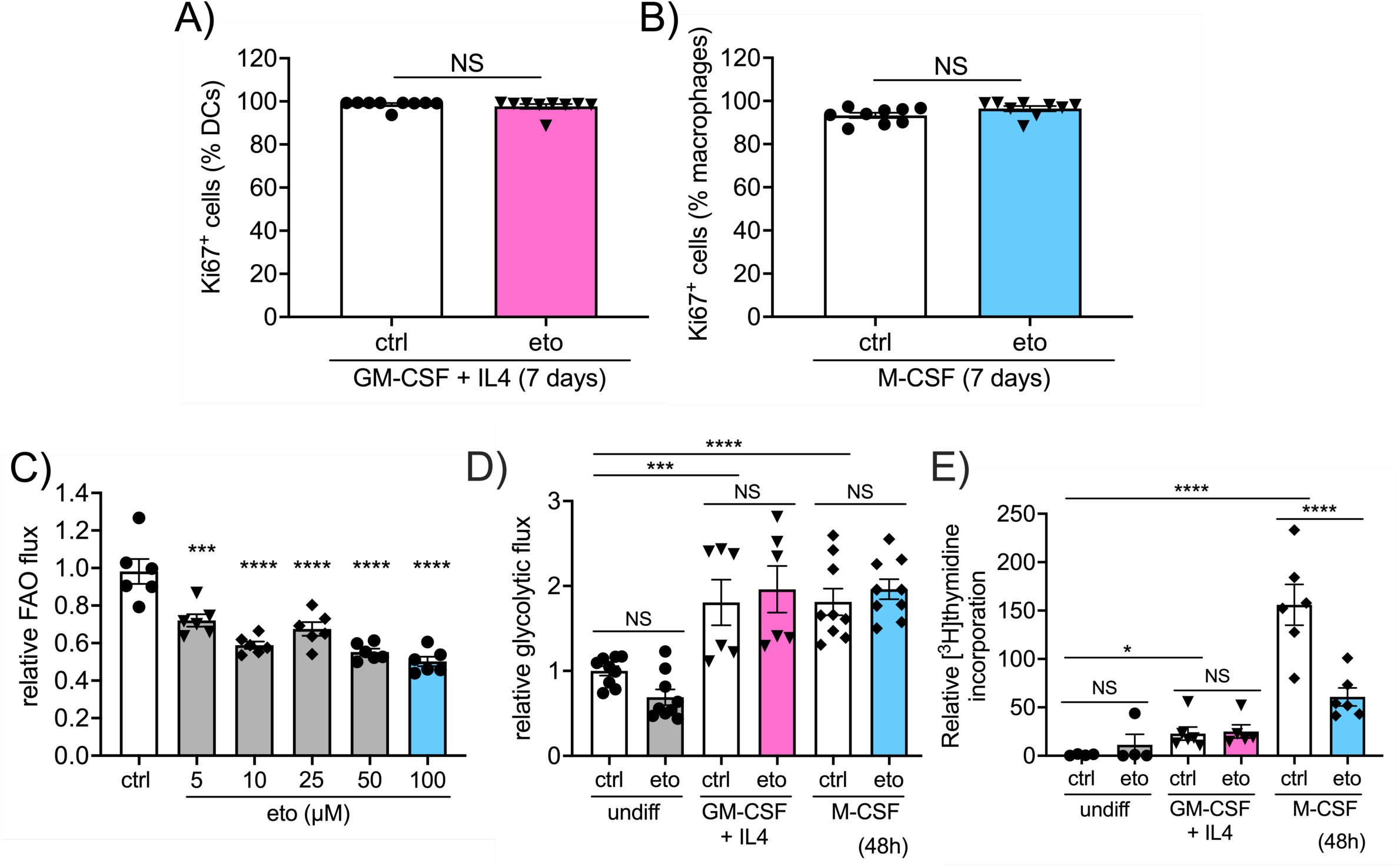
Assessment of FAO inhibition on proliferation and glycolytic flux from in vitro monocyte differentiation. **A-F)** Data from in vitro differentiation of bone marrow-derived monocytes stimulated towards DC (GM-CSF + IL4) or macrophage (M-CSF) lineages. **A**,**B)** Flow cytometric assessment of cells after 7 days culture. **A)** Proliferating DCs as assessed by Ki67 as a percentage CD11c^+^MHCII^+^CD64^−^ cells (ctrl, n=9; eto, n=9). **B)** Proliferating macrophages assessed by Ki67 as a percentage of CD11b^+^CD64^+^CD24^−^ cells (ctrl, n=9; eto, n=9). **C)** FAO flux in monocytes treated with 5, 10, 25, 50 or 100 µM etomoxir for 24h in basal media, as assessed by 9,10-[^3^H]palmitic acid radioisotope incorporation (n=6). **D)** Glycolytic flux in monocytes in media only, or treated for 48h with GM-CSF + IL4 or M-CSF, as assessed by [5-^3^H]glucose radioisotopic incorporation (ctrl, n=6; eto, n=6). **E)** Proliferation of monocytes in media only, or treated for 48h with GM-CSF + IL4 or M-CSF, as assessed by [^3^H]thymidine incorporation (ctrl, n=6; eto, n=6). Data are represented as mean of individual data points from at 3 independent experiments ± SEM. NS, not statistically significant; \**p*<0.05; \*\*\**p*<0.001; \*\*\*\**p*<0.0001; NS, not statistically significant, by *t*-test **(A**,**B)** or ANOVA and Bonferroni post-hoc test **(C-E)**.

**Supplemental Figure 4:**
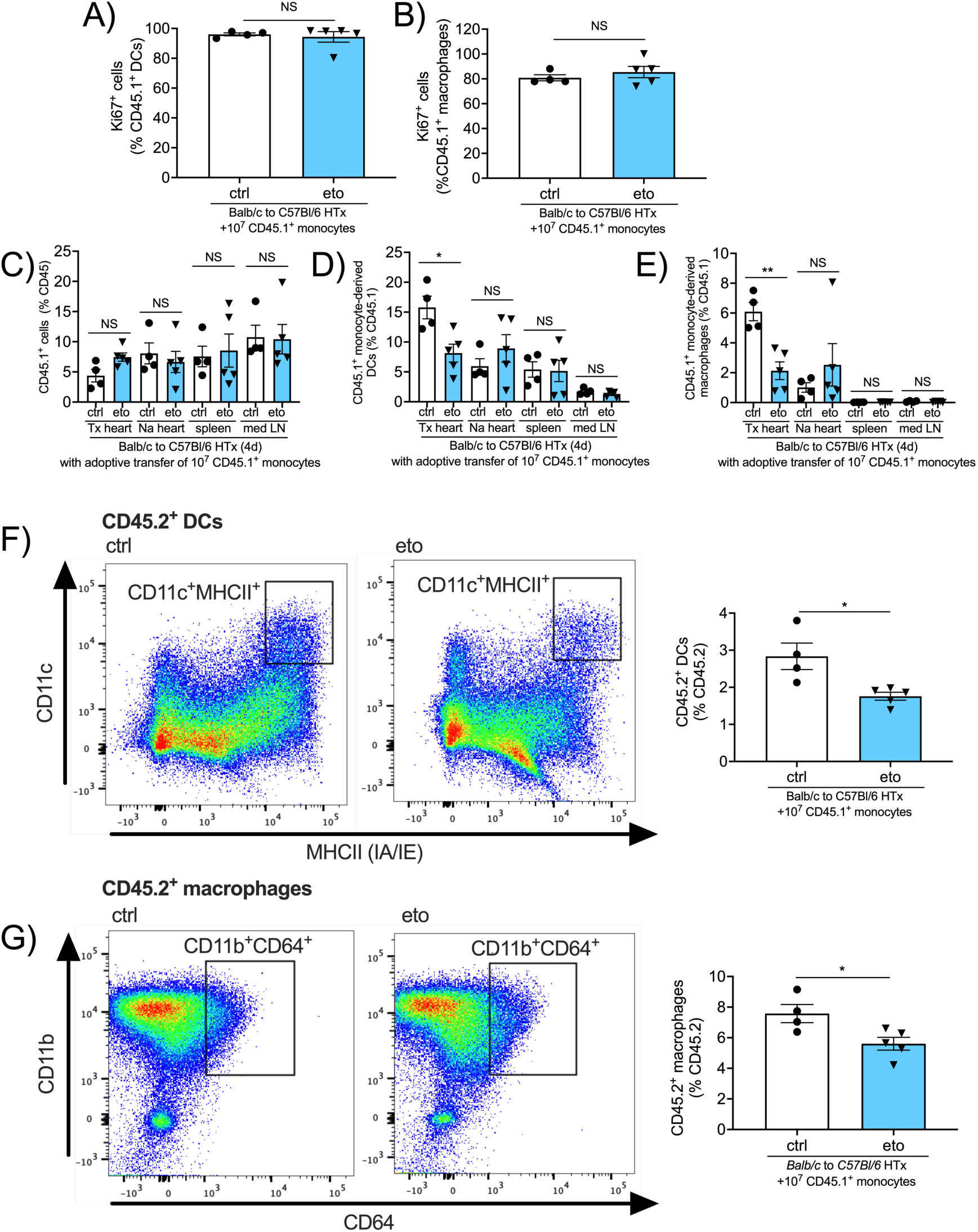
Assessment of CD45.1^+^ monocytes and monocyte, DC and macrophage proliferation in adoptive transfer experiments. **A-J)** Flow cytometric assessment of transplanted hearts at 4 days post-HTx from Balb → B6 HTx adoptively transferred with CD45.1^+^ monocytes. **A**,**B)** Assessment of proliferation by percentage Ki67 incorporation. **A)** In CD11c^+^MHCII^+^CD64^−^ dendritic cells (DCs) as a percentage of CD45.1^+^ cells (ctrl, n=4; eto, n=5). **B)** In CD11b^+^CD64^+^CD24^−^ macrophages as a percentage of CD45.1^+^ (ctrl, n=4; eto, n=5). CD45.1^+^ cells as a percentage of CD45^+^ cells in the transplanted heart (Tx heart), native heart (Na heart), spleen and mediastinal lymph node (med LN) (ctrl, n=4; eto, n=5). **D)** CD11c^+^MHCII^+^CD64^−^ dendritic cells (DCs) as a percentage of CD45.1^+^ cells in the Tx heart, Na heart, spleen and med LN (ctrl, n=4; eto, n=5). **E)** CD11b^+^CD64^+^CD24^−^ macrophages as a percentage of CD45.1^+^ cells in the Tx heart, Na heart, spleen and med LN (ctrl, n=4; eto, n=5). **F)** CD45.2^+^ DCs assessed by CD11c^+^MHCII^+^CD64^−^ cells (ctrl, n=4; eto, n=5). **G)** CD45.2^+^ macrophages assessed by CD11b^+^CD64^+^CD24^−^ cells (ctrl, n=4; eto, n=5). Data are represented as mean ± SEM. NS, not statistically significant; \**p*<0.05; NS, not statistically significant, by *t*-test (**A**,**B, F**,**G**) or ANOVA and Bonferroni post-hoc test (**C-E**).

**Supplemental Figure 5:**
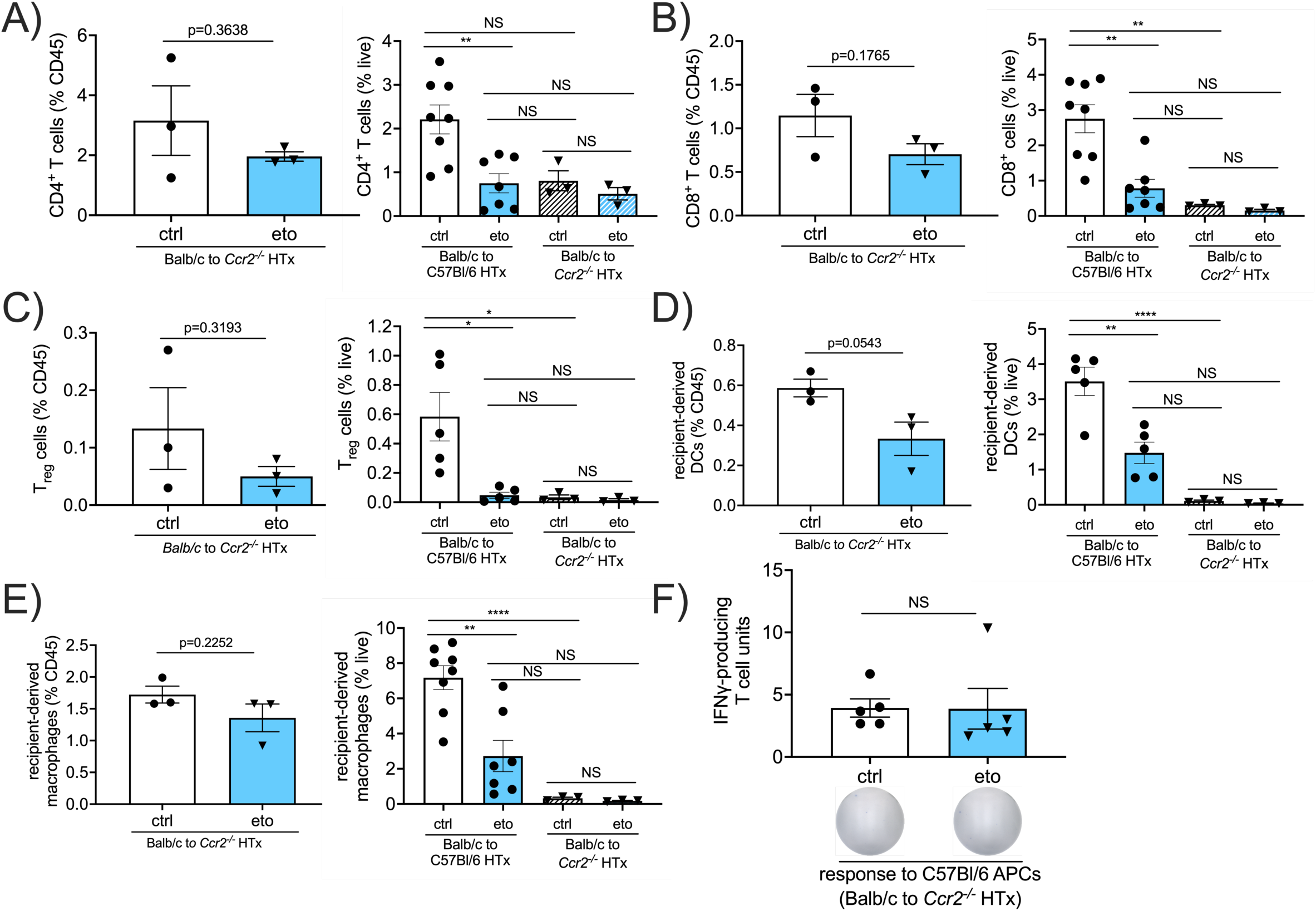
Flow cytometry for T cells in *C**cr**2*^*-/-*^ HTx. **A-E)** Flow cytometric assessment in transplanted hearts from Balb → *Ccr2*^*-/-*^heart allografts at 4 days post-transplant shown as a percentage of CD45^+^ cells (left graph) or live cells (right graph; data for live cells from Balb → B6 HTx is re-plotted from Figs. 1 and 2). **A)** CD4 T cells assessed by CD4^+^CD3^+^CD90.2^+^ cells (ctrl, n=8; eto=7 for Balb → B6 HTx; ctrl, n=3; eto, n=3 for Balb → B6 *Ccr2*^*-/-*^HTx). **B)** CD8 T cells assessed by CD8^+^CD3^+^CD90.2^+^ cells (ctrl, n=8; eto=7 for Balb → B6 HTx; ctrl, n=3; eto, n=3 for Balb → B6 *Ccr2*^*-/-*^HTx). **C)** T regulatory cells assessed by Foxp3^+^CD4^+^CD3^+^CD90.2^+^ cells (ctrl, n=8; eto=7 for Balb → B6 HTx; ctrl, n=3; eto, n=3 for Balb → B6 *Ccr2*^*-/-*^HTx). **D)** Recipient-derived dendritic cells assessed by IA-b^+^CD11c^+^MHCII^+^CD64^−^ cells (ctrl, n=5; eto=5 for Balb → B6 HTx; ctrl, n=3; eto, n=3 for Balb → B6 *Ccr2*^*-/-*^HTx). **E)** Recipient-derived macrophages assessed by IA-b^+^CD11b^+^CD64^+^CD24^−^ cells (ctrl, n=8; eto=7 for Balb → B6 HTx; ctrl, n=3; eto, n=3 for Balb → B6 *Ccr2*^*-/-*^HTx). **F)** ELISPOT assay performed using splenocytes procured at 4d post-transplantation from Balb → *Ccr2*^*-/-*^heart allografts from either vehicle (ctrl) or etomoxir (eto)-treated recipients. Samples were stimulated with activated B6 APCs and assessed for the production of IFNγ (ctrl, n=4; eto, n=4). Data are represented as mean ± SEM. NS, not statistically significant; \*\**p*<0.01; \*\*\**p*<0.001; \*\*\*\**p*<0.0001 by *t*-test (**A-E left panels; F**) or ANOVA and Bonferroni post-hoc test (**A-E right panels**).

**Supplemental Figure 6:**
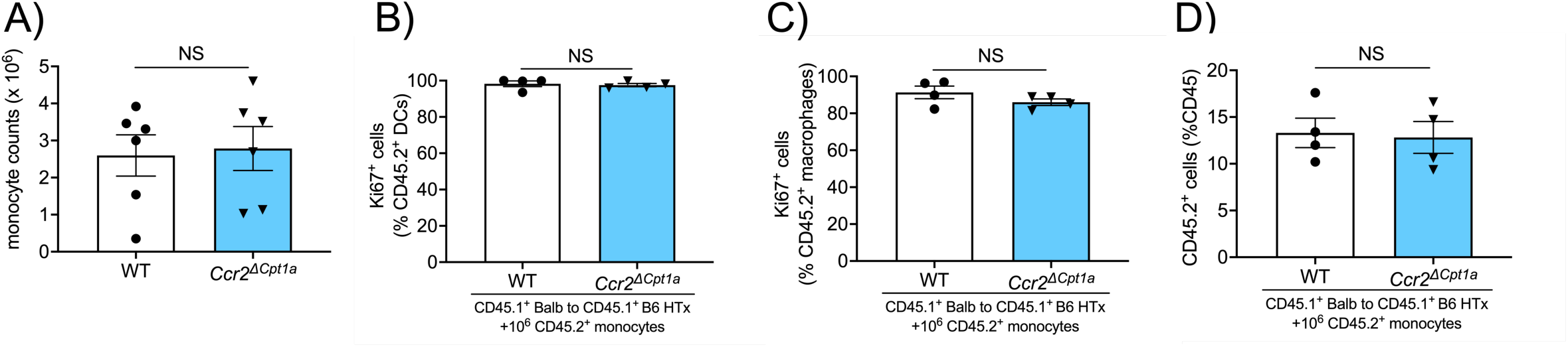
Genetic deletion of Cpt1a on proliferation of monocyte-derived macrophages and monocyte-derived DCs in heart allografts. **A)** Cell counts after monocyte isolation from the bone marrow of *Cre-negative;Cpt1a*^*fl/fl*^ (WT; n=6) or *Ccr2*.*Cre*^*ER*^*;Cpt1a*^*fl/fl*^ (*Ccr2*^*ΔCpt1a*^; n=6) mice. **B-C)** Assessment of proliferation by percentage Ki67 incorporation in CD45.2^+^ fraction from transplanted hearts in Balb to B6 CD45.1^+^ HTx recipients adoptively transferred with WT or *Ccr2*^*ΔCpt1a*^ monocytes. **B)** In CD11c^+^MHCII^+^CD64^−^ dendritic cells (DCs) as a percentage of CD45.2^+^ cells (WT, n=4; *Ccr2*^*ΔCpt1a*^, n=4). **C)** In CD11b^+^CD64^+^CD24^−^ macrophages as a percentage of CD45.2^+^ (WT, n=4; *Ccr2*^*ΔCpt1a*^, n=4). **D)** In CD45.2^+^ cells as a percentage of CD45^+^ cells (WT, n=4; *Ccr2*^*ΔCpt1a*^, n=4). Data are represented as mean ± SEM. \**p*<0.05; NS, not statistically significant, by *t*-test.

## Tables

**Supplemental Table 1.** List of antibodies used for flow cytometry.

| antibody | clone | company |
| --- | --- | --- |
| CD11b | M1/70 | eBioscience (San Diego, CA) |
| CD3ε | 17A2 | eBioscience |
| CD4 | GK1.5 | eBioscience |
| CD45R (B220) | RA3-6B2 | eBioscience |
| CD64 | X54-5/7.1 | eBioscience |
| CD90.2 (Thy-1.2) | 30-12 | eBioscience |
| Ki-67 | SolA15 | eBioscience |
| Ly-6G | 1A8 | eBioscience |
| MHC class II (I-A/I-E) | 114.15.2 | eBioscience |
| NK1.1 | PK136 | eBioscience |
| CD11b | M1/70 | BD Biosciences (San Jose, CA) |
| CD11c | N418 | BD Biosciences |
| CD11c | HL3 | BD Biosciences |
| CD115 | T38-320 | BD Biosciences |
| CD24 | M1/69 | BD Biosciences |
| CD3ε | 145-2C11 | BD Biosciences |
| CD45 | 30-F11 | BD Biosciences |
| CD45.1 | A20 | BD Biosciences |
| CD45.2 | 104 | BD Biosciences |
| CD64 | X54-5/7.1 | BD Biosciences |
| CD69 | H1.2F3 | BD Biosciences |
| CD8a | 53-6.7 | BD Biosciences |
| Foxp3 | M23 | BD Biosciences |
| I-A[b] | AF-1201 | BD Biosciences |
| Ly-6G | 1A8 | BD Biosciences |
| MHC class II (I-A/I-E) | M5/114.15.2 | BD Biosciences |
| NK1.1 | PK136 | BD Biosciences |
| CD11c | N418 | Biolegend (San Diego, CA) |
| Foxp3 | FJK-16s | Thermo Scientific (Waltham, MA) |

